# Systematic imaging reveals features and changing localization of mRNAs in *Drosophila* development

**DOI:** 10.1101/008938

**Authors:** Helena Jambor, Surendranath Vineeth, Alex T. Kalinka, Pavel Mejstrik, Stephan Saalfeld, Pavel Tomancak

## Abstract

mRNA localization is critical for eukaryotic cells and affects numerous transcripts, yet how cells regulate distribution of many mRNAs to their subcellular destinations is still unknown. We combined transcriptomics and systematic imaging to determine tissue-specific expression and subcellular distribution of 5862 mRNAs during *Drosophila* oogenesis. mRNA localization is widespread in the ovary, depends on the microtubule cytoskeleton and the mRNAs enriched at the posterior cortex share common localization machinery. Localized mRNAs, in particular the posterior class, have distinct gene features and differ in expression level, 3’UTR length and sequence conservation from unlocalized mRNAs. Using cross-tissue comparison we revealed that the localization status of mRNAs differs between epithelial, germline and embryonic cell types and also changes within one cell, the oocyte, over time. This dataset enables the transition from deep mechanistic dissection of singular mRNA localization events towards global understanding of how mRNAs transcribed in the nucleus distribute in cells.

## Introduction

Cell differentiation is accompanied by polarization and segregation of membranes, cytoplasm and organelles. A powerful mechanism to generate subcellular asymmetries used by eukaryotes and even prokaryotes is mRNA localization in combination with controlled protein translation (reviewed in Medioni et al., 2012). Long-range mRNA transport in most metazoans relies on the polarized cytoskeleton and the microtubule minus-and plus-end motor complexes. mRNA enrichment at microtubule minus-ends is aberrant in mutants that affect the dynein motor complex, while plus-end directed transport requires kinesin molecules (reviewed in Medioni et al., 2012, Bullock, 2011).

Mechanistic dissection of the canonical localisation examples showed that, mRNAs localize through cis-regulatory sequences, zipcodes, which are often present in the 3’UTR of the transcript (reviewed in Jambhekar and Derisi, 2007) and zipcode-binding proteins that initiate the formation of transport competent ribonucleoproteins (RNPs) (Chao et al., 2010, Dienstbier et al., 2009, Bullock et al., 2010, Dix et al., 2013). mRNAs can also harbor two antagonizing localization signals that act consecutively in cells and direct mRNAs sequentially to opposing microtubule ends (Ghosh et al., 2012; Jambor et al., 2014), suggesting that transport RNPs could be regulated. It has further been shown that some mRNA localization elements are active in several cell types suggesting that the mRNA transport machinery is widely expressed and mRNA localization elements function in a cell-type independent manner (Bullock and Ish-Horowicz, 2001, Jambor et al., 2014, Snee et al., 2005, Kislauskis et al., 1994).

In addition to microtubule-based transport, few mRNAs can enrich by trapping to a localized anchoring activity (Forrest and Gavis, 2003, Sinsimer et al., 2011) or by hitch-hiking along with a localization-competent mRNA (Jambor et al., 2011). Recent live-imaging revealed that the same mRNA can, depending on the cell type use both diffusion and active transport mechanism (Park et al., 2014). Further *in vitro* data show that mRNA transport along microtubules can occur both uni-and bi-directional, suggesting mRNAs can switch between processive and diffusive transport modes (Soundararajan and Bullock, 2014).

mRNA localization is perhaps best characterized in the oocyte of *Drosophila melanogaster (D.melanogaster)* where localization of *oskar, bicoid* and *gurken* is instrumental for setting up the embryonic axes (Neuman-Silberberg and Schüpbach, 1993, Ephrussi et al., 1991, Berleth et al., 1988, St Johnston et al., 1989). However, more recent work suggests that mRNA localization is not occurring only for few singular mRNAs but instead is a widespread cellular feature that affects a large proportion of expressed mRNAs {Shepard, 2003; Blower, 2007; Lecuyer, 2007; Zivraj, 2010; Cajigas, 2012). How a cell distinguishes localized from ubiquitous transcripts and orchestrates transport of many mRNAs remains enigmatic. It is conceivable that each localised mRNAs carries its own zipcode sequence that directs it to specific subcellular location. However, despite wealth of data on co-localised transcripts computational methods thus far fail to detect such signals in a reliable manner. Alternatively co-packaging of several mRNA species, only one of which carries specific localisation signal, has been shown in at least two cases (Jambor et al., 2011, Lange et al., 2008) but seems to be less common in neuronal and embryonic cell types (Mikl et al., 2011, Amrute-Nayak and Bullock, 2012). It is also unclear whether the mRNA localization status differs between cell types and to what extent is it subject to tissue specific regulation.

Here we unravel the global landscape of mRNA localization in the *Drosophila* ovary by combining stage specific mRNA sequencing with genome-wide fluorescent in situ hybridizations (FISH) and systematic imaging. The localized transcripts show characteristic gene level features such as longer and highly conserved 3’UTRs that clearly distinguish subcellular enriched and ubiquitous mRNAs and is most pronounced in the class of posterior localized mRNAs. Cross-comparison with previous datasets on RNA localisation in embryos (Lecuyer et al., 2007) revealed that mRNA localization diverges across cell types and within one cell over time. These changes are not due to alternative transcript expression since the germline cells of the Drosophila ovary show only little transcriptional change. While all subcellular mRNAs tested require an intact microtubule cytoskeleton for their transport, posterior mRNAs further depend on initially localized oskar mRNA.

## Results

### Widespread mRNA localization in ovaries

We systematically probed and imaged the expression and subcellular distributions of mRNAs in *Drosophila* mass-isolated egg chambers followed by stage-specific mRNA sequencing (3Pseq and RNAseq) and genome-wide fluorescent in situ hybridization (FISH). RNA sequencing data, expression pattern annotations (using a hierarchical controlled vocabulary^1^) and images (representative 2D images and original z-stacks) are collected in a publicly accessible database, the Dresden Ovary Table, DOT^2^ (Figure 1- figure supplement 1A, B). We identified 3624 mRNAs as being expressed based on our in situ hybridization screen, most of which were also detectable in the two RNAseq datasets that are in good agreement with each other (Figure 1A, Figure 1-figure supplement 2A, Figure 2-figure supplement 2A). 65% of these were expressed ubiquitously in ovarian cells at all time-points of oogenesis (ubiquitous), but we also observed expression in subsets of cells (13%, cellular) and mRNAs that asymmetrically localized in the cytoplasm (22%, subcellular) or the nucleus (5%, nuclear).

**Figure 1.**
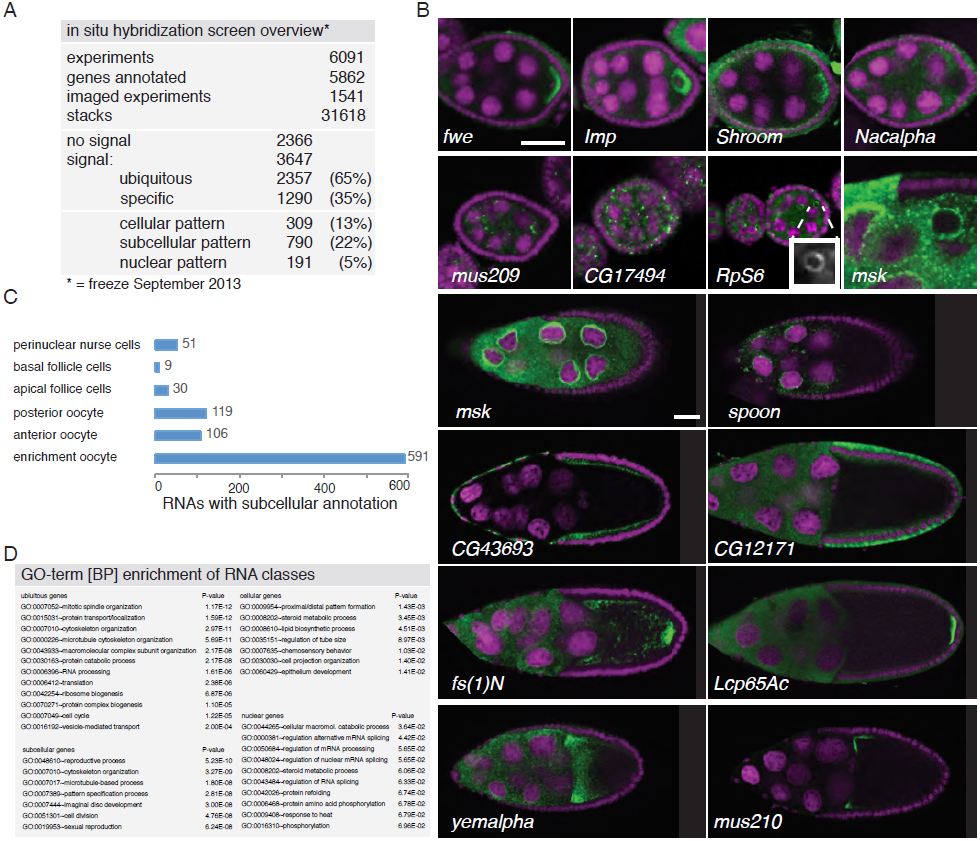
Summary of the fluorescent in situ hybridization (FISH) screen on ovaries.

**(A)**. Summary of key numbers of the screen. For each of the 6091 FISH experiment we annotated the signal as “no signal”, “ubiquitous” or “specific”. Only specific and some ubiquitous signals were imaged. **(B)**. Exemplary subcel-lular expression patterns. In the syncytial early egg-chamber, 590 mRNAs are transported from the site of transcription in the nurse cells into the developing oocyte. Here mRNAs are either restricted to a cortical domain (fwe) or detectable in the entire ooplasm (Imp). Occasionally mRNAs simultaneously enriched in the oocyte portion of the syncytial egg-chamber and at the apical membrane of the somatic epithelial cells (Shroom). Five mRNAs were specifically excluded from the oocyte portion and enriched in the nurse cells (Nacalpha). Few mRNAs were enriched anterior in stage 2-7 oocytes (mus209). mRNAs showed ubiquitous granules in the cytoplasm (CG17494) or rarely ubiquitous ring-like staining patterns (RpS6, arrow; 10x10μm inset showing only the RNA channel. mRNAs also enriched around the nucleus of the oocyte (msk) and the nurse cells nuclei (msk) and this varied from an entire ring around the nucleus to specific sub-areas of the perinuclear space (spoon). Apical enrichment was detected in late epithelial somatic cells (CG43693) while basal localization in the follicle cells was relatively rare (CG12171). Anterior and posterior RNA localization varied between diffuse (fs(1)N, yemalpha) and tight cortical enrichments (Lcp65Ac, mus210). **(C)**. Distribution of subcellular localized mRNAs in subcategories. Note: mRNAs can appear in more than one subgroup. **(D)**. GO-term enrichment analysis of ubiquitous, cellular, nuclear and subcellular gene sets.

The 309 mRNAs of the cellular category were predominantly expressed in the somatic epithelium and often restricted to subsets of cells and specific time-points (Figure 1- figure supplement 2B, C). 191 RNAs were detectable specifically in ovarian nuclei mostly of the endocycling, polyploid nurse cells, but also in epithelial cells and in 29 cases in the oocyte nucleus (Figure 1-figure supplement 2D,E). Oocyte nuclear mRNAs showed temporal changes and could be nurse cell transcripts imported into the oocyte nucleus or instances of transcription from the meiotic nucleus (Saunders and Cohen, 1999). Nuclear patterns varied from ring-like signal to dispersed foci or widespread distribution in the nucleoplasm and were not linked to the chromosomal position of the genes (Figure 1- figure supplement 2F). Precursors of micro RNAs and long non-coding RNAs also showed varying degrees of nuclear enrichments (Figure 1-figure supplement 2G).

Subcellular mRNA localization affected 790 mRNAs, but was limited to small number of subcellular domains (Figure 1B, C). The largest group, 591 mRNAs, was enrichment in the oocyte portion of the syncytial egg-chamber during early oogenesis (*fwe, Imp, Shroom*) where also microtubule minus-ends are concentrated. At mid-oogenesis the oocyte establishes its own polarized microtubule cytoskeleton; at this stage, we observed 106 mRNAs enriched towards the anterior and 119 mRNAs enriched at the posterior pole, corresponding to where the microtubule minus and plus ends are enriched (Theurkauf et al., 1992, Januschke et al., 2006). The quality of these localizations ranged from tight (*mus210, Lcp65Ac*) to diffuse (*yemalpha, fs(1)N*) association at the anterior-dorsal, the entire anterior or the posterior cortex. mRNAs were also detected in subcellular domains of the nurse (*msk, spoon*) and epithelial cells (*CG43693, CG12171*). For few mRNAs, we observed previously unknown ovary accumulations for example mRNAs in cytoplasmic granules (*CG17494*), depleted from the oocyte (*Nacalpha*), showing cortical enrichment (*Actn*) or forming ring-like structures (*CG14639,* Figure 1B, Figure 1-figure supplement 2H).

We next asked whether the ubiquitous, nuclear, cellular and subcellular gene sets are functionally related. Gene Ontology (GO) analysis showed that subcellular mRNAs are enriched for terms describing reproductive processes, cytoskeleton organization and cell cycle regulation, while the cellular gene set, being mostly expressed in the somatic cells, was enriched for GO-terms describing epithelial development, lipid trafficking and cuticle formation and nuclear RNAs enriched for terms assigning them to RNA regulatory processes (Figure 1D). These four categories also maintain distinct expression patterns during embryogenesis. Genes of the subcellular set are enriched among genes expressed in the central nervous system, suggesting an interesting relatedness of these polarized tissues (Figure 1-figure supplement 3).

### Changing mRNA localization during development

Since most ovarian genes are expressed again during embryogenesis in distinct patterns, we next wondered whether also subcellular localization is preserved in different cell types during development. We took advantage of the wealth of FISH data now available for *Drosophila* and combined our data for the somatic epithelial cells and the germline cells of the ovary with the FISH screen performed on embryonic cells (Lecuyer et al., 2007). These screens in combination covered 9114 genes of which 1674 mRNAs showed subcellular localization at least at one time-point either during oogenesis or embryogenesis and thus are “localization competent” (Figure 2-figure supplement 1A,B). Filtering of the datasets for mRNAs that were probed by FISH in all three cell types resulted in 720 mRNAs of which only five mRNAs were localized in all three, and 89 mRNAs were localized in two cell types (Figure 2A). Since mRNA localization to the microtubule minus-ends in somatic epithelia, germline and embryonic cells is mechanistically equivalent (Bullock and Ish-Horowicz, 2001, Jambor et al., 2014), we asked whether the same mRNAs tend to localize to the same microtubule end in different cell types. The data show that only three mRNAs were localized to microtubule minus end in all three cell types (*Dok, Sdc, CG12006;* Figure 2A). Other localization classes had similarly little overlap across cell types (Figure 2-figure supplement 1C). Thus we conclude that while many mRNAs are localization competent, these localizations appear to be transient and mRNAs are generally not constitutively localized during development.

**Figure 2.**
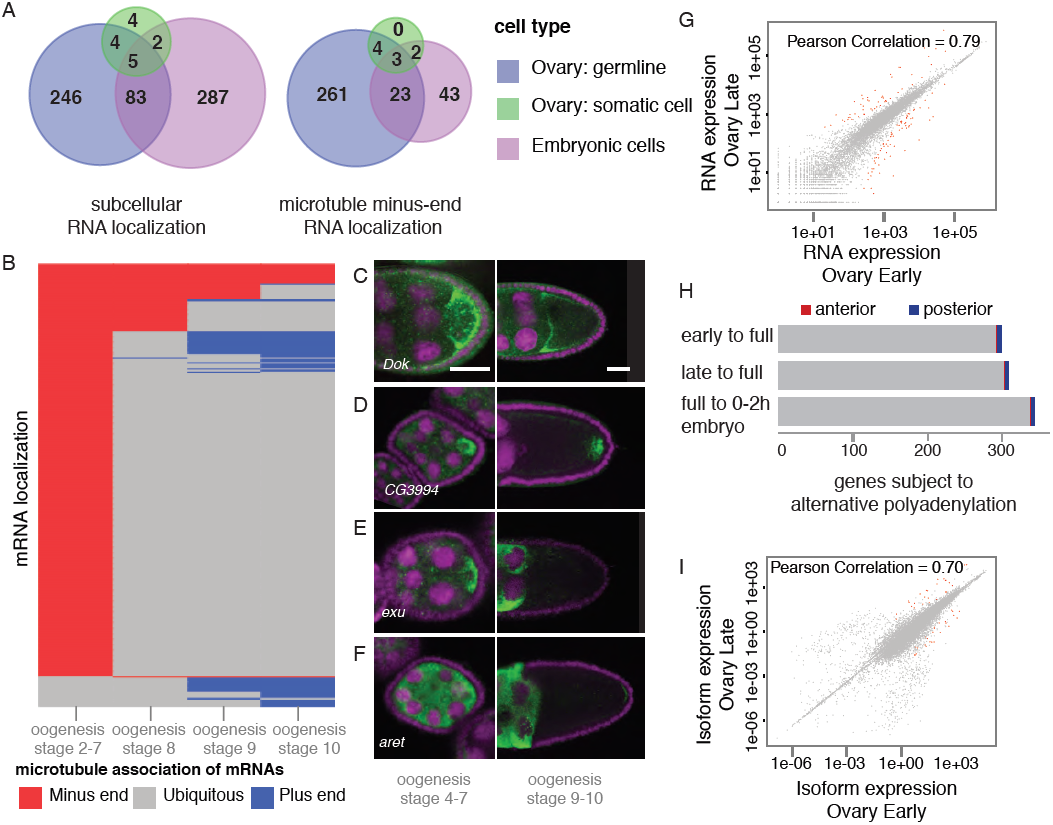
mRNA localizations are highly variable across tissues and within cells. **(A)**. Venn-diagrams globally comparing mRNA localization (left) and microtubule minus-end co-localization (right) in somatic, germline and embryonic cell types. Only 5 (<1%) mRNAs localized in all sampled cell types, 89 (14%) mRNAs localized in at least two cell types. Similarly, only 3 mRNAs (<1%) constitutively co-localized with microtubule minus-ends, localization in two cell types occurred for 29 mRNAs (9%). **(B-F)**. mRNA localizations in one cell, the oocyte, over time. **(B)**. Dendrogram where each line represents the localization of single mRNAs in the oocyte from early to late oogenesis; indicated are microtubule minus-end co-localization (red), plus-end co-localization (blue) and ubiquitous phases of mRNA distribution (grey). **(C)**. Dok mRNA remains localized at minus-ends from early to stage 10 of oogenesis. **(D)**. CG3994 mRNA initially co-localizes with microtubule minus-ends but switches to a plus-end accumulation at stage 9/10 of oogensis. **(E)**. After an initial microtubule minus-end co-localization exu mRNA becomes ubiquitously distributed. **(F)**. The initially ubiquitously distributed aret mRNA adopt a weak plus-end accumulation at stage 9. **(C-F)**. FISH showing the RNA in green and DNA (labelled with DAPI) in magenta. Scale bar 30μm. **(G)**. Pair-wise correlation of early/late 3Pseq data revealed that the stage-specific transcriptomes were highly similar (Pearson Correlation: 0.79); only few genes, highlighted in red, were significantly up-or down-regulated (p-value adjusted for multiple testing < 0.1). **(H)**. Only ∼300 genes (early-full: 298; late-full: 308; full-embryo: 346) changed their mean-weighted 3’UTR length that is indicative of an alternative polyadenylation. Alternative UTR form usage was found for 1 to 4 anterior (red) and 4 to 5 posterior (blue) mRNAs during oogenesis. **(I)**. Correlation analysis of expressed transcript isoform (deduced from RNAseq data) revealed that from early to late ovaries almost no transcript-isoforms significantly changed in their expression level. Transcripts with significant changes are shown in red.

To address whether mRNA localization is more stable within one cell type, we compared the oocyte-localized mRNAs at different time-points of oogenesis up to the onset of embryonic transcription. Clustering mRNAs by microtubule association revealed that even within one cell mRNA localization is not a stable property. Only 22% of the mRNAs were localized constitutively (Figure 2B) and the majority changed their specific subcellular destinations over time. During early oogenesis, the majority of mRNAs (n=591) co-localized with microtubule minus ends, however less than 100 mRNAs remained localized at later stages. Of those, 39 remained associated with microtubule minus ends (Figure 2C) while 66 changed to microtubule plus-end co-localization either at stage 9 or at stage 10 (Figure 2D). Most minus-end transcripts however switched to a ubiquitous state at mid-oogenesis (Figure 2E). mRNAs could also adopt de-novo plus-end localization at stage 9 and 10 after having been ubiquitously distributed during early oogenesis (Figure 2B, F). Such de-novo accumulation of an initially ubiquitous transcript was not observed at microtubule minus-ends.

Lastly, also the global dynamics of changes in mRNA localization differed: the minus-end mRNAs rapidly de-localized (stage 8: n=99; stage 10: n=39) while in contrast plus-end localized mRNAs continuously increased (stage 8: n=68; stage 10: n=109) and this was further accentuated when including early embryogenesis (Figure 2-figure supplement 1D). Minus-end accumulation only re-emerged after initiation of zygotic transcription of the embryo, pointing to a potential link between transcription and minus-end localization of mRNAs, analogous to the known link between nuclear events and microtubule plus-end localization (Hachet and Ephrussi, 2004, Besse et al., 2009).

The surprising changes in mRNA localization status across cell types and within one cell over time prompted us to ask whether this could be explained by transcriptional regulation during oogenesis. Alternative splicing was previously shown to differentially regulate mRNA localization by producing localized and non-localized isoforms of the same gene (Whittaker et al., 1999, Horne-Badovinac and Bilder, 2008). We therefore probed our stage-specific transcriptomic data for changes in gene expression. In agreement with results from gene expression analyses of whole ovaries measured by microarray (Chintapalli et al., 2007) and RNAseq (Graveley et al., 2011) we find that about half of the *D. melanogaster* genes were expressed at each sampled time point and the vast majority of these expressed transcripts, 85%, were detectable at every time point from early oogenesis until embryogenesis (Figure 2-figure supplement 2B,C). Further the expression levels across time points were highly correlated (Figure 2G, Figure 2-figure supplement 2D), suggesting that the transcriptome remained constant throughout oogenesis. A significant up-or down regulation of gene expression levels was only observed for 626 transcripts and among them are only rare examples of germline specific transcripts (padj< 0.1, Figure 2G: red data points, Supplementary Table 2-4, Figure 2- figure supplement 2E,F). Instead, GO-term analysis associated genes under differential expression with extracellular matrix, vitelline membrane and cuticle formation, consistent with their expression in the somatic epithelial cells (Figure 2-figure supplement 2E).

Across the entire oogenesis, we also could not detect shortening or lengthening of the 3’UTRs, changes in the number of transcript ends and while 55% of genes were expressed in alternative isoforms, the vast majority (>99%) of genes showed no change in isoform expression (Figure 2H,I, Figure 2-figure supplement 2G, H-J, Supplementary Table 5-7). Furthermore, the ubiquitous gene set showed similar transcript diversity as subcellular genes. Thus, expression of different mRNA variants, cannot account for the many changes in mRNA localization status across cell types and within the oocyte over time. The little variation in mRNA expression starting from egg chamber formation until the onset of zygotic transcription suggests that oogenesis is not dependent on transcriptional changes but rather on post-transcriptional regulation of the expressed transcripts, in particular through mRNA localization.

### Posterior localizations depend on *oskar* mRNA

The time-course analysis of mRNA distributions in the oocyte showed large groups of transcripts that recapitulate the localization pattern of the well-characterized, singular mRNAs such as *oskar, gurken, nanos* and *bicoid.* This suggests that these mRNAs each are representatives for an mRNA localization class that could be co-regulated. Transport of mRNAs towards the anterior and the posterior pole of the oocyte requires an intact microtubule cytoskeleton; accordingly, the localization of all new anterior and posterior candidate mRNAs is lost in colchicine treated egg-chambers, while ubiquitously distributed mRNAs or RNA foci in the nucleoplasm, that lacks a microtubule cytoskeleton, were unaffected by the colchicine treatment (Figure 3A, Figure 3-figure supplement 1A-E, Supplementary Table 8).

**Figure 3.**
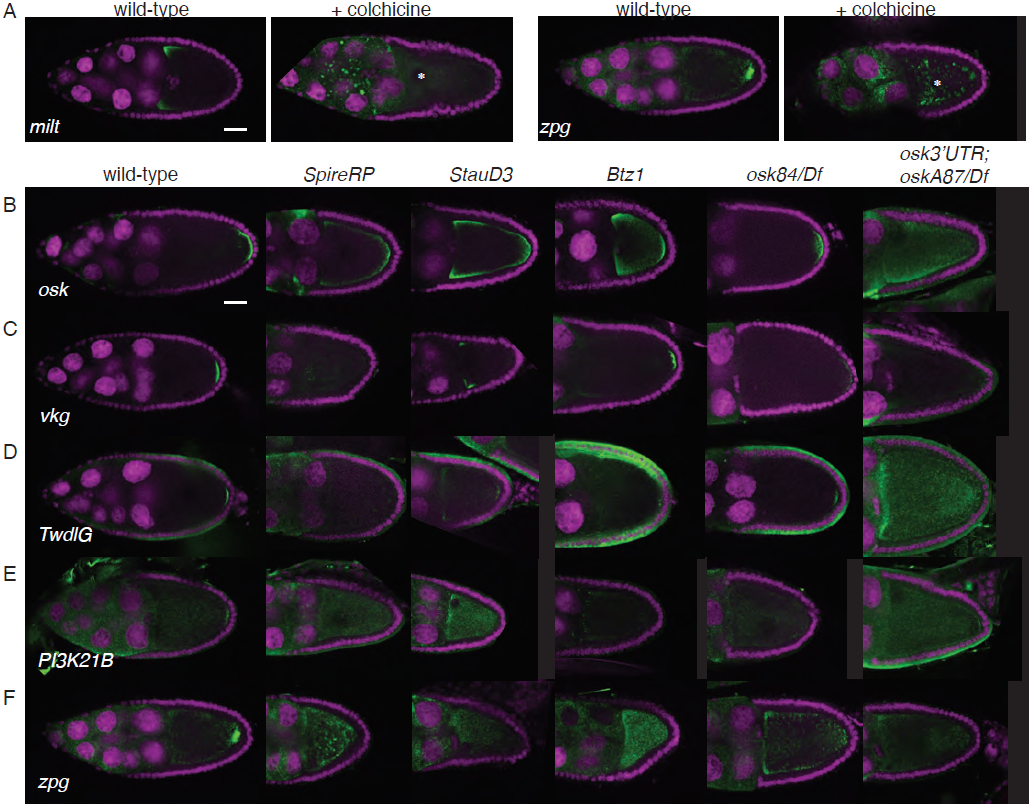
mRNA localization requires the microtubule cytoskeleton and posterior enrichment is impaired in posterior localization pathway mutants. **(A-F)**. FISH experiments showing the RNA in green and DNA (labelled with DAPI) in magenta. Scale bars 30μm. **(A)**. Localization of an exemplary anterior (*milt*) and posterior mRNA (*zpg*) is lost upon microtubule depolymerization by colchicine. See summary of the results in Figure S6E and accompanying Supplementary Table 8. **(B-F)**. Localization of the novel posterior candidate mRNAs *vkg* **(C)**, *TwdlG* **(D)**, *PI3K21B* **(E)** and *zpg* **(F)** is lost in egg-chambers that prematurely depolymerize the microtubules (flies homozygous for *SpireRP),* are mutant for the RNA binding protein Staufen (flies homozygous for *StauD3*) or mutant for the EJC protein Barentz (flies homozygous for *Btz1*). The candidate mRNAs are mis-localized in a manner similar to *oskar* mRNA **(B)**, whose localization is known to be disrupted in those mutant conditions. In *Btz1* egg chambers a weak enrichment of *vkg* mRNA remained that in rare instances is also observed for *oskar* mRNA. The localization of the tested novel posterior mRNAs was also lost at stage 10 in egg-chambers mutant for Oskar protein (*osk84/Df(3R)pXT103*). All candidate mRNAs were lost in egg-chambers that do not express posterior *oskar* mRNA. Egg-chambers used were from *oskar* RNA null flies that express the *oskar* 3’UTR in order to rescue the early oogenesis arrest (*oskar 3’UTR/+; oskA87/Df(3R)pXT103;* Jenny et al., 2006).

To compare in more detail the localization mechanism of new candidates we focused on the group of posterior localized transcripts that show localization similar to the known posterior mRNA, *oskar.* Posterior transport of *oskar* mRNA requires components of the EJC complex, the RNA binding protein Staufen and an intact microtubule cytoskeleton while maintenance of *oskar* localization beyond stage 9 needs Oskar protein to anchor the mRNA (Hachet and Ephrussi, 2004, van Eeden et al., 2001, St Johnston et al., 1991, Ephrussi et al., 1991, Micklem et al., 2000, Vanzo and Ephrussi, 2002). The posterior enrichment of the selected candidate mRNAs was severely reduced in egg-chambers mutant for EJC components (*Btz*^*1*^), that have a disrupted cytoskeleton (*Spire*^*RP*^), lack Staufen (*Stau*^*D3*^) and Oskar protein (*osk*^*84*^/*Df(3R)p*^*XT103*^) and strongly resembled the mis-localized *oskar* mRNA (Figure 3B-F). Thus the novel posterior mRNAs require the same proteins for their localization as *oskar* mRNA.

To investigate whether the novel candidate mRNAs use the posterior transport machinery independently of *oskar* mRNA we used a genetic combination in which egg-chambers lack any localized *oskar* mRNA at the posterior cortex (Jenny et al., 2006). None of the new posterior mRNAs showed posterior localization in these egg-chambers (Figure 3B-F, Figure 3-figure supplement 1F), indicating that the novel posterior mRNAs depend on *oskar* mRNA for their localization. This *oskar* mRNA-dependent localization could either be due to the role of *oskar* in recruiting and stabilizing microtubule plus-ends at the posterior pole (Zimyanin et al., 2007) or *oskar* mRNA could serve as a platform that allows mRNA to hitch-hike to the posterior pole (Jambor et al., 2011), possibly in a large transport granule. The latter possibility could suggest that not all posterior mRNAs have their own posterior-zipcode for their individual transport. Instead, posterior transport could involve co-packaging via RNA-RNA or RNA-protein interaction motifs.

### Global features of localized mRNAs

The dynamic mRNA localizations in oocytes and the *oskar* mRNA dependency for localization of posterior candidate mRNAs suggest a more multifaceted regulation of mRNA transport than through zipcode-recognition alone. We therefore next asked whether there are global gene features that could set localized mRNAs apart from ubiquitous ones. Ovarian expressed mRNAs differed in their expression levels over several orders of magnitude. Using our stage specific 3Pseq data we analysed the expression levels for each gene set. Ubiquitous and subcellular mRNA expression levels were overall comparable however, the posterior class was significantly higher expressed than all other localization classes, including the related anterior mRNAs (Figure 4A, Figure 4-figure supplement 1A). Considering how seemingly inefficient posterior transport is (Zimyanin et al., 2008), higher expression levels could be an additional measure to ensure that enough mRNAs will eventually localize. In particular the late phase accumulation of posterior localized mRNAs in the enlarged oocyte (Sinsimer et al., 2011, Forrest and Gavis, 2003) could benefit from high expression levels.

**Figure 4.**
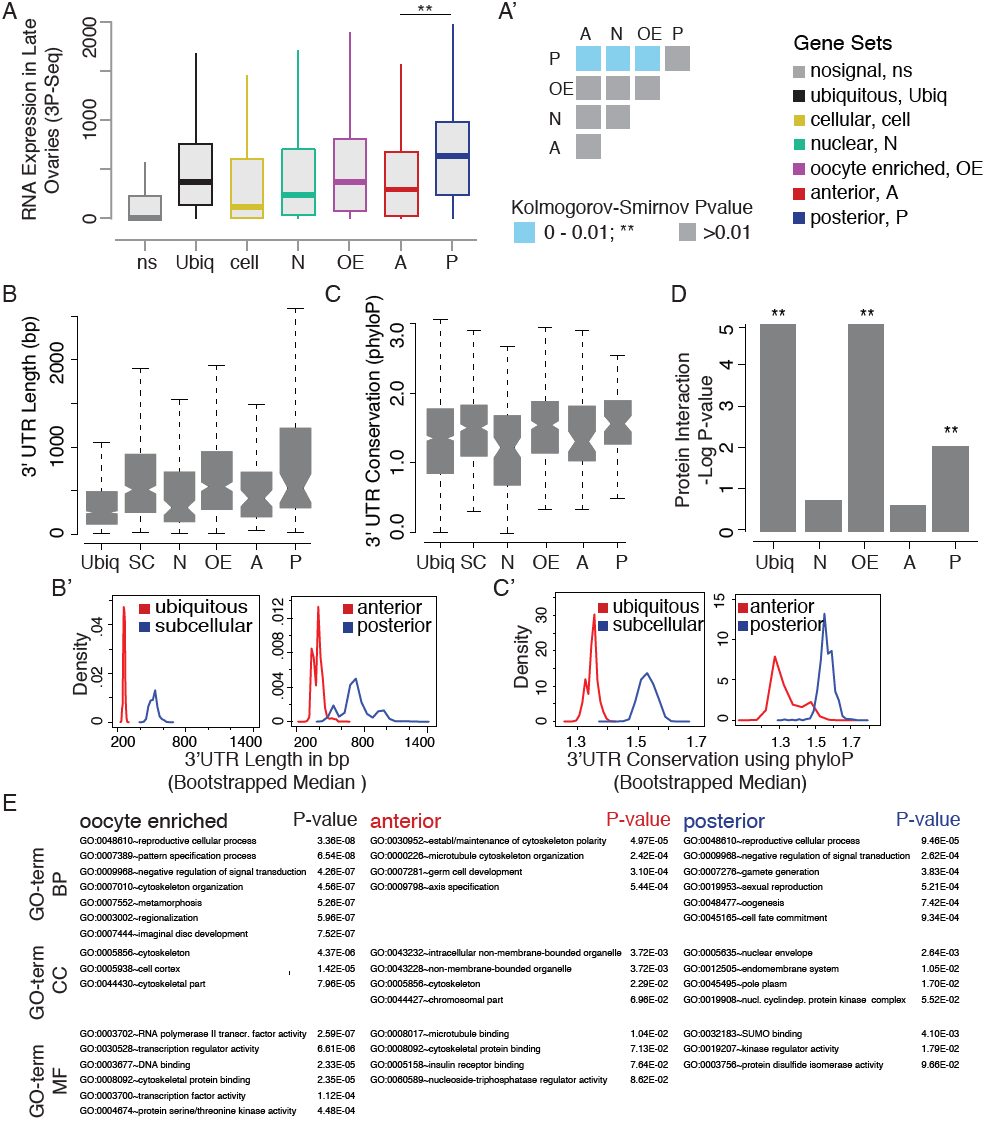
Subclasses of localized mRNAs and their specific features at the level of gene architecture, mRNA expression, evolutionary conservation and function. **(A)**. Boxplots showing distributions of median mRNA expression levels for gene sets defined by annotations (see Supple-mentary Table 1). Shown are 3Pseq quantifications from late ovary mRNA (for early, full ovaries and early embryogenesis see Figure S5A). mRNAs of the posterior group showed significantly higher expression than anterior mRNAs (a’, Kolmogorov-Smirnov p-value: 3.9e-06). **(B)**. Distributions of 3’UTR length for gene sets. **(B’)**. Results of a non-parametric randomization test to show that ubiquitous and subcellular genes (p-value = 0) and anterior and posterior genes (p-value = 0.0018) have significantly different median 3’UTR lengths (i.e. no or little overlap of densities). SC = subcellular gene set. **(C)**. Median conservation of the 3’UTR sequence for gene sets across 24 Drosophila species. **(C’)**. Result of a non-parametric randomization test showing that ubiquitous genes are significantly less conserved in their 3’UTRs than subcellu-lar genes (p-value: 0) and posterior genes show higher conservation than anterior genes (p-value: 0.0032). SC = subcellular gene set. **(D)**. Protein interaction analysis per gene set revealed that posterior genes, but not anterior genes, share significantly more protein-protein interactions than would be expected by chance. **(E)**. GO-terms associated with oocyte enriched, anterior and posterior gene sets. Shown are the p-values for each GO-term calculated by the modified Fisher Exact test, which results in the EASE score p-value.

Yet, expression level alone cannot account for subcellular localization. We therefore compared the gene-level variables of each localization class and revealed that subcellular mRNAs had significantly longer 3’UTR sequences and this was more pronounced for the posterior localization class (Figure 4B,B’). The posterior gene set further showed longer gene structures, longer 5’UTRs, longer exons and introns, a higher number of exons and introns and a higher intron proportion compared to ubiquitous and anterior mRNAs (Figure 4-figure supplement 1B-H). Consistent with the observation that localized mRNAs are enriched in non-coding portions, the exon proportion was highest in the ubiquitous gene set (Figure 4-figure supplement 1I). The high intron proportion of posterior genes is particularly interesting in light of the recent finding that the stable deposition of the EJC, required for posterior *oskar* mRNA localization, is correlated with long intron-containing genes (Ghosh et al., 2012, Ashton-Beaucage et al., 2010). Localized genes not only had longer 3’UTRs, but also showed higher 3’UTR sequence conservation than ubiquitous genes, and again this was significantly more pronounced in the posterior gene set (Figure 4C, C’). Longer 3’UTRs could harbour several conserved motifs that in sum lead to the changing subcellular mRNA enrichments across time.

Analysis of protein interaction data shows that proteins of the posterior gene set participate in significantly more protein-protein interactions, suggesting that the close proximity of their transcripts in the cell could be of functional importance (Figure 4D). To gain further insight into potential biological functions of mRNA localization classes, we performed GO-enrichment analysis for oocyte enriched, anterior and posterior gene sets that linked all categories to reproductive and patterning processes (Figure 4E). Anterior and oocyte enriched mRNAs were further enriched for terms describing cytoskeleton regulation, which also was overrepresented in all “localization competent” mRNAs (Figure 2-figure supplement 1B). However, while the oocyte enriched class associated with microtubule and actin cytoskeleton terms, the anterior gene set only enriched for microtubule terms. The observation that localized mRNAs are highly enriched for cytoskeletal regulators is particularly interesting in light of a recent model suggesting a self-organizing principle for the polarized cytoskeleton in mouse neurites (Preitner et al., 2014). Anterior mRNAs are additionally associated with chromosome and cell cycle regulation. Taken together, the close proximity of anterior localized mRNAs with the oocyte nucleus suggests a potential role for mRNA localization in female meiosis. In contrast, the posterior mRNAs associated strongly with signalling, cell fate commitment and membrane organization terms. This is consistent with the known signalling events between the oocyte and the overlaying somatic epithelial cells at the posterior pole (Roth et al., 1995) and with the membrane re-modelling as the germ plasm is being assembled (Vanzo et al., 2007, Tanaka and Nakamura, 2008).

### Discussion

To conclude, we generated a comprehensive resource, the Dresden Ovary Table (DOT) that includes stage-specific RNA expression and subcellular localization data for the entire oogenesis from cystoblast division to the beginning of embryogenesis. This genome-wide approach allowed us to define gene sets of co-localized mRNAs and show that localized, particularly posterior mRNAs, have a more complex gene structure, longer and higher conserved non-coding features and higher expression levels than ubiquitous mRNAs.

Our resource consists of 52,000 carefully selected, annotated, stage specific images of ovarian gene expression and localization patterns that can be searched online or downloaded for in-depth computational analysis. The curated images are linked to 32,000 raw 3D image stacks available for interactive browsing that will facilitate further discovery by the scientific community. The ovary dataset is integrated with similar data on gene expression and RNA localization patterns in *Drosophila* embryos (Tomancak et al. 2007, Lecuyer et al. 2007) enabling comparisons between tissues on a gene-by-gene basis.

Cross-tissue and time-course analyses revealed the changing mRNA localization profile during development and that the well-described, canonical examples of mRNA localization in the ovary (Neuman-Silberberg and Schüpbach, 1993, Ephrussi et al., 1991, Berleth et al., 1988, St Johnston et al., 1989) represent classes of co-regulated mRNAs. The extent of the changes in localization status during development was however unexpected, especially considering that during oogenesis the transcriptome appears stable. The pervasive changes in localization status of mRNAs are in stark contrast to the observations that mRNAs localize through sequence encoded mRNA zipcodes (reviewed in Medioni et al., 2012) and that the localization machinery is active in all cell types analysed (Jambor et al., 2014, Bullock and Ish-Horowicz, 2001). It will therefore be interesting to ask whether specificity of mRNA localization is based on selective, cell-type specific mRNA regulation machinery or a zipcode signal that is under specific temporal control. mRNAs can for example harbour two consecutively acting localization signals that direct mRNAs sequentially to opposing microtubule ends (Ghosh et al., 2012, Jambor et al., 2014). However, how one signal is de-activated and the other activated is yet unknown. Alternatively, only few mRNAs could have a zipcode for their localization and the vast majority would be co-transported with these regulated mRNAs in large transport granules. Finally, it is also conceivable that subcellular mRNAs could be locally trapped by unidentified physical properties of subcellular cytoplasmic domains. Our observation that all examined posterior mRNAs fail to localize in the absence of *oskar* mRNAs lends support to the idea that these mRNAs are hitchhiking the *oskar* localization machinery and might not themselves have a “posterior”-zipcode. Regardless of the specific mechanisms of mRNA transport, our genome-wide analysis shows that mRNA localization is a phenomenon contingent on the cellular context and is most likely highly regulated during development. Our dataset enables the transition from deep mechanistic dissection of singular RNA localization events towards systemic examination of how RNAs transcribed in the nucleus distribute in cells and how this affects cellular architecture and cell behaviour in development.

## Materials and Methods

### Mass isolation of *Drosophila* egg-chambers

Flies were grown under standard laboratory conditions, fed for 2 days with fresh yeast at 21 and 25 °C. For isolation of egg-chambers we developed a mass-solation protocol (see below) that allows us to enrich separated egg-chambers of all stages.

### RNA isolation, sequencing and analysis

We isolated total mRNA using TRIreagent (Sigma Aldrich) from stage 1-7 egg-chambers, including the germline stem cells, from stage 8-10 egg-chambers and from total ovaries containing mainly stage 11 and older egg-chambers. Additionally, RNA from 0-2h embryos was isolated. We used two complementary mRNA sequencing approaches; standard whole mRNA sequencing (RNAseq) and a sequencing method that captures specifically the sequence adjacent to the poly(A) tail and thus allows direct counting of transcripts (3Pseq, V. Surendranath and A. Dahl, manuscript in preparation). Of the ∼50 million (3Pseq) and 100 million (RNAseq) Illumina reads we mapped 70% (3Pseq) and 90% (RNAseq) to the *D.melanogaster* release 5.52 genome with Bowtie. Quantification was done using HTSeq (Anders and Huber, 2010). Normalization and differential expression was done using DESeq (Anders and Huber, 2010). Noise thresholds of 70 and 50 counts, for RNAseq and 3Pseq respectively, were derived from observing the distributions of normalized counts. 3’ UTR forms were assigned by overlaying annotated Flybase UTR forms with 3Pseq reads lying within 200 nucleotides of the annotated 3’ UTR end. Alternate Polyadenylation events were called by calculating the mean weighted UTR length (Ulitsky et al., 2012), a difference of 200 nucleotides in the mean weighted lengths corresponding to two biological stages resulted in the gene being considered as undergoing Alternate Polyadenylation.

### 96-well Fluorescent in situ Hybridization

We used an established protocol for in situ hybridization in 96-well plates (Tomancak et al., 2007) with minor adaptations (see below): we added an over-night wash step after hybridization, incubate the anti-DIG antibody over night and used fluorescent tyramides for probe detection. Each experiment was evaluated and imaged using a wide-field microscope (Zeiss Axioplan Imaging) equipped with an optical sectioning device (DSD1, Andor) to generate confocal-like z-stacks.

### Annotation and Database

We developed a controlled vocabulary to describe the cell types and relevant subcellular structures for oogenesis for germline and somatic cells^3^. Experiments showing no detectable FISH signal were classified as “no signal at all stages”, while experiments resulting in a homogeneous signal throughout oogenesis were classified as “ubiquitous signal at all stages”. Gene expression patterns were imaged up to stage 10B of oogenesis after which cuticle deposition prevents probe penetration. Each pattern that did not fall in the above-mentioned classes was imaged at all stages of oogenesis in several individual egg-chambers per time-point. We collected 3D images and used custom scripts in FIJI (Schindelin et al., 2012) to manually select and orient representative 2D images that were uploaded to the Dresden Ovarian-expression Table^4^ (DOT). The 2D images remain linked to the original image stacks and all the raw stacks that were used to create an exemplary 2D image are available for interactive inspection using a simple image browsing cgi script. Thus the record of each in situ experiment for a given gene consists of a set of 2D images assigned to a specific oogenesis stage and described using annotation terms selected from the controlled vocabulary. For definition of broad classifications, subclass grouping and embryo annotation class definition see Supplementary Table 1.

### Binary matrix

The binary matrix summarizes the data of our screen in tabular form, which facilitates access to the multidimensional image annotation data and integrates them with the RNAseq and 3Pseq data. The binary matrix is a freeze from September 2013, based on which our analyses were done. The binary matrix is provided as a flat file for independent bioinformatics investigation of the dataset^5^.

The matrix contains the following information for each annotated gene:

the FlyBase ID; the expression levels as raw as well as normalized counts from RNAseq and 3Pseq experiments for early -, late - and full ovaries and 0-2h embryos; the pair-wise comparison of expression over the time course analysed, raw and normalized; the mean-weighted length indicating alternative 3’UTR expression.

The binary matrix additionally contains the annotation of FISH expression patterns. The expression terms are from the controlled vocabulary (CV). If the CV term is true its value is equal to 1 otherwise it is zero. If a gene is annotated twice during the screen, the CV values are summed up and thus result in values >1.

We also provide information which clone was used to prepare the FISH probe; the classification into broad annotation classes (“no signal”; “ubiquitous”; “specific”, see Results), classification of specific expression patterns into subclasses (“cellular”, “subcellular”, “nuclear”); reliability status: “reliable” and “non-reliable” (genes probed with more than one RNA probe that resulted in conflicting annotations (n=247), were labelled as “not reliable”. 185 “unreliable” cases resulted from a “no signal” versus “ubiquitous” or “no signal” versus “specific” annotations, here we assume one of the probes to be non-functional. 57 “unreliable” annotations were due to different probes giving a “ubiquitous” and “specific” signal respectively. One possibility is that probes were specific to different isoforms of the gene.); pn-status: comparison of sequencing and FISH results (TN = true negative: genes expressed below cut-off in either RNAseq or 3Pseq and giving a “no signal” in FISH experiments. FN = false negatives: genes expressed below cut-off in either RNAseq or 3Pseq and giving a “ubiquitous” or “specific” in FISH signal. TP = true positives: genes expressed above cut-off in either RNAseq or 3Pseq and giving a “ubiquitous” or “specific” in FISH signal. FP = false positives: genes expressed above cut-off in either RNAseq or 3Pseq and resulting in a “no signal” FISH annotation (see Figure 1-figure supplement 2A).

### GO-term analysis

For GO-term enrichment of gene sets we used the DAVID web server (Huang da et al., 2009). Terms or features enriched at a false discovery rate (FDR) of ≤10% and/or a Benjamini p-value of <0.1 were considered significant. Two stringencies were applied: the standard FDR cut-off (≤10%) or the more stringent ‘Benjamini’ p-value (≤0.1).

### Colchicine treatment and Mutant analysis

Flies were fed for 15 hours at 25°C with fresh yeast paste supplemented with 50μg/ml colchicine (Cha et al., 2002). The effect of colchicine on individual egg-chambers was determined by scoring the detachment of the oocyte nucleus from the anterior cortex and its migration towards the centre of the oocyte. To test posterior localization in mutants that affect *oskar* mRNA localization we used ovaries from homozygous *Spire*^*RP*^ (Manseau and Schupbach, 1989), *Stau*^*D3*^ (St Johnston et al., 1991) and *Btz*^*1*^ (van Eeden et al., 2001) flies. Further we analysed egg-chambers from *osk84/* Df(3R)p^XT103^ flies lacking functional Oskar protein (Lehmann and Nusslein-Volhard, 1986) and from oskar3’UTR/+;oskA87/ Df(3R)p^XT103^ flies that entirely lack endogenous *oskar* mRNA but develop past the early oogenesis arrest characteristic for *oskar* RNA null flies due to a transgenic source of oskar 3’UTR (Jenny et al., 2006) that is incapable posterior localization.

### Gene feature variable

For analysis of the annotated gene features we used the flybase gff data (*D.melanogaster* release 5.52) processed using custom Python scripts.

### 3’UTR length and conservation

For each gene, we defined the most used UTR form as the form that was most highly expressed (relative to any other forms expressed from the same gene) and which had UTR ends that overlapped by +/-200 bp with a FlyBase annotated UTR end. From this data, we extracted unique 3’ UTR lengths for each gene. Sequence conservation of 3’ UTRs was measured as median phyloP scores (Pollard et al., 2010) across all bases in the most used UTR form for 3’ UTR sequence alignments across 24 *Drosophila* species (using the *D. melanogaster* UTR co-ordinates). PhyloP scores were calculated using the R package *Rphast* (Hubisz et al., 2011). Median UTR lengths and conservation scores were bootstrapped by re-sampling genes with replacement from selected annotation sets 100,000 times and calculating median values for each re-sample. p-values were calculated as the number of re-samples in which the annotation group with a lower median value was greater than or equal to the re-sampled median of the annotation group to which it was being compared, divided by 100,000.

### Protein Interactions

A manually-curated *D. melanogaster* protein-protein interaction network was downloaded from the mentha interactome database (Calderone et al., 2013). To test whether genes belonging to certain annotation groups participated in more protein-protein interactions within the annotation group than expected by chance, we adopted the following randomization-based approach. A random sample, the size of the number of genes in an annotation group that participate in at least one interaction in the total protein interactome, was taken from the total set of genes belonging to the protein interactome, and the number of protein interactions within this random sample was scored, minus loops. This was repeated 100,000 times to generate a distribution of the number of interactions obtained by randomly sampling the number of genes belonging to the annotation group from the total interactome. The p-value was calculated as the number of randomly sampled networks that had as many or more interactions as the real annotation group divided by 100,000.

### Protocol: Mass-isolation of egg-chambers

1. Flies were fed with fresh yeast and kept for 1-2 days at 25°C.
2. Mixed sex flies were narcotized with CO_2_ for a maximum of 5 minutes before proceeding to step 3.
3. Narcotized flies were immediately immersed in 4% Formaldehyde in PBS (for FISH experiments) or in PBS supplemented with 0.1% Tween-10 (for ovarian extract or total RNA isolation).
4. Flies were rapidly processed twice through a grinding mill adaptor at a fine setting (grade step “3”) on a standard food processor (Kitchen Aid).
5. The ground flies were size-separated using 850, 450 and 212μm sieves successively, resulting in a flow-through highly enriched for separated egg-chambers of all stages.
6. Collection of mass-isolated material:

a. For FISH experiments the co-isolation of testis and gut materials did not disturb the subsequent analysis and the material was allowed to settle by gravity and to be fixed for additional 15 minutes in 4% Formaldehyde, resulting in an overall fixation time of 20 minutes. The supernatant was then removed, the material washed twice in 1xPBS and then transferred stepwise into 100% methanol for storage at -20°C.
b. For isolation of total RNA we manually selected egg-chambers at early stages (germarium to stage 7, previtellogenesis), late stages (stage 9-10, postvitellogenesis) and full ovaries highly enriched for stage 11+ egg-chambers using a stereomicroscope. For each stage we collected at least 10μ1 of total material that was frozen immediately.

### Protocol: 96-well plate fluorescent in situ hybridization (FISH)

1. Mass isolated egg-chambers were transferred stepwise (MeOH/PBT 3:1; MeOH/PBS 1:1; MeOH/PBS 1:3) into PBT0.1% (each wash few minutes)
2. Egg-chambers were then washed 6x in PBT0.1%, 5 minutes each
3. Egg-chambers were briefly washed in PBT0.1%/Hyb 1:1
4. Pre-Hybridization of egg-chambers was done in 200μ1 hybridization buffer at 55 °C for 1 hour.
5. Egg-chambers were then added to a 96-well plate and hybridized over-night at 55°C in Hybridization Buffer *with* Dextran Sulfate supplemented with 2μ1 of probe.
6. 100μl of warm Wash Buffer was added to each well and immediately removed together with probe-solution.
7. Egg-chambers were rinsed once with 150μl of Wash Buffer and then washed four times for one hour at 55°C in Wash Buffer.
8. Egg-chambers were then washed five times for 1hr at 55 °C in 150μ1 PBT, the last wash was done over-night at 55°C.
9. Egg-chambers were washed twice for 1hr at room temperature in 150μ1 PBT.
10. The primary antibody (Anti-Digoxigenin-POD Fab Fragments (Roche) was diluted 1:200 and egg-chambers were incubated in 200μ1 antibody solution over-nmRNA expression, evoluight.
11. Egg-chambers were rinsed with 150μ1 of PBT and then washed ten times for 30 minutes at RT in 150μ1 of PBT0.1%.
12. For detection egg-chambers were incubated with Cy3-Tyramides (Perkin-Elmer) 1:70 diluted in amplification buffer for 30 minutes.
13. Egg-chambers were then washed ten times for 30 minutes at room temperature in 150μ1 of PBT. DAPI, diluted 1:1000, was included in one wash step.
14. All PBT was removed and ∼50μ1 mounting medium was added.

## Footnotes

1 http://tomancak-srv1.mpi-cbg.de/cgi-bin/ovary_annotation_hierarchy.pl

2 http://tomancak-srv1.mpi-cbg.de/DOT/main

3 http://tomancak-srv1.mpi-cbg.de/cgi-bin/ovary_annotation_hierarchy.pl

4 http://tomancak-srv1.mpi-cbg.de/DOT/main

5 http://tomancak-srv1.mpi-cbg.de/cgi-bin/dump_binary_matrix_ovary.pl?db=insitu_ovaries

## Acknowledgement

We thank Diana Selig, Jens Schmiedel and David Seniuk for image acquisition and processing, Holger Brandl for bioinformatic services, Franziska Friedrich for drawings of egg-chambers, Anne Starkloff for webpage, Andreas Dahl, Deep Sequencing Group at CRTD/BIOTECH, Dresden for generation of the 3Pseq data, Anne Ephrussi and Daniel St Johnston for fly lines, Michael Hiller for sharing the 24 *Drosophila* species alignment. We are grateful for discussions of the manuscript to Florence Besse, Simon Bullock, Carsten Hoege, James Saenz and Vitaly Zimyanin. V.S. received support from DIGS-BB. H.J. and P.T. were supported by FP7-EU. Project: GENCODYS. P.T. was additionally funded by HFSP Young Investigator Grant RGY0093/2012 and by The European Research Council Community′s Seventh Framework Program (FP7/2007-2013) grant agreement 260746.

## Author contribution

H.J. designed and performed the experiments, participated in the bioinformatics analysis, and wrote the manuscript. V.S. designed and performed most bioinformatic analyses including mapping, quantification of RNAseq and 3’seq reads, and statistical analysis of gene architecture features. A.T.K. performed statistical analysis of 3’UTR length, evolutionary conservation, and protein interactome data. P.M. performed the 96-well in situ hybridization. S.S. designed an image processing script to enhance analysis of fluorescent image data. P.T. co-designed the study, performed cross-tissue analysis, developed database and website, and co-wrote the manuscript.

## Author information

The RNAseq data is deposited under the SRA accession number is SRP045258.

The authors declare no competing financial interests.

---

**Figure 1-figure supplement 1.**
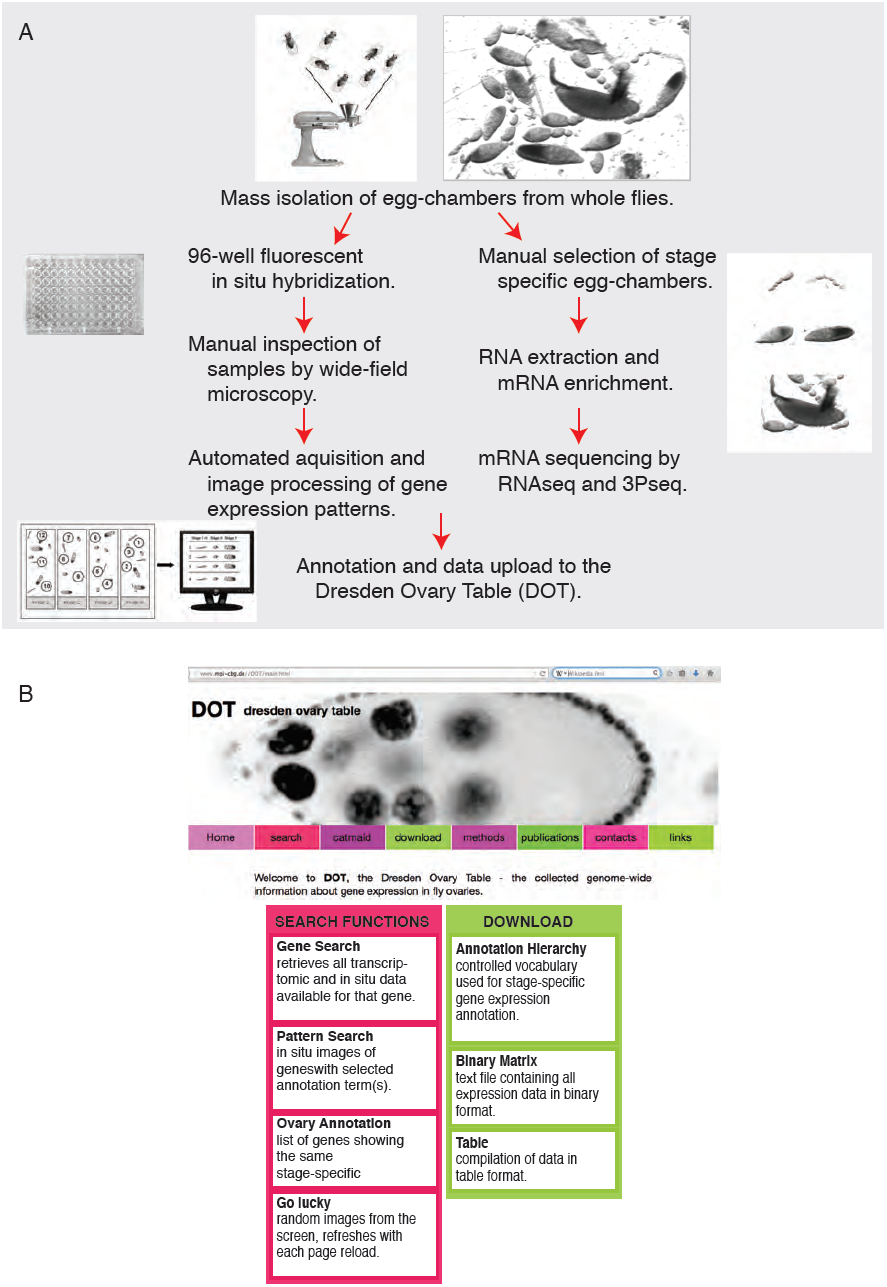
Experimental outline and database features. **(A)**. Overview of the experimental procedure for transcriptome and genome-wide in situ hybridization experiments and evaluation. **(B)**. Screenshot of the publicly available Dresden ovary table, DOT, and key search and download functions.

**Figure 1-figure supplement 2.**
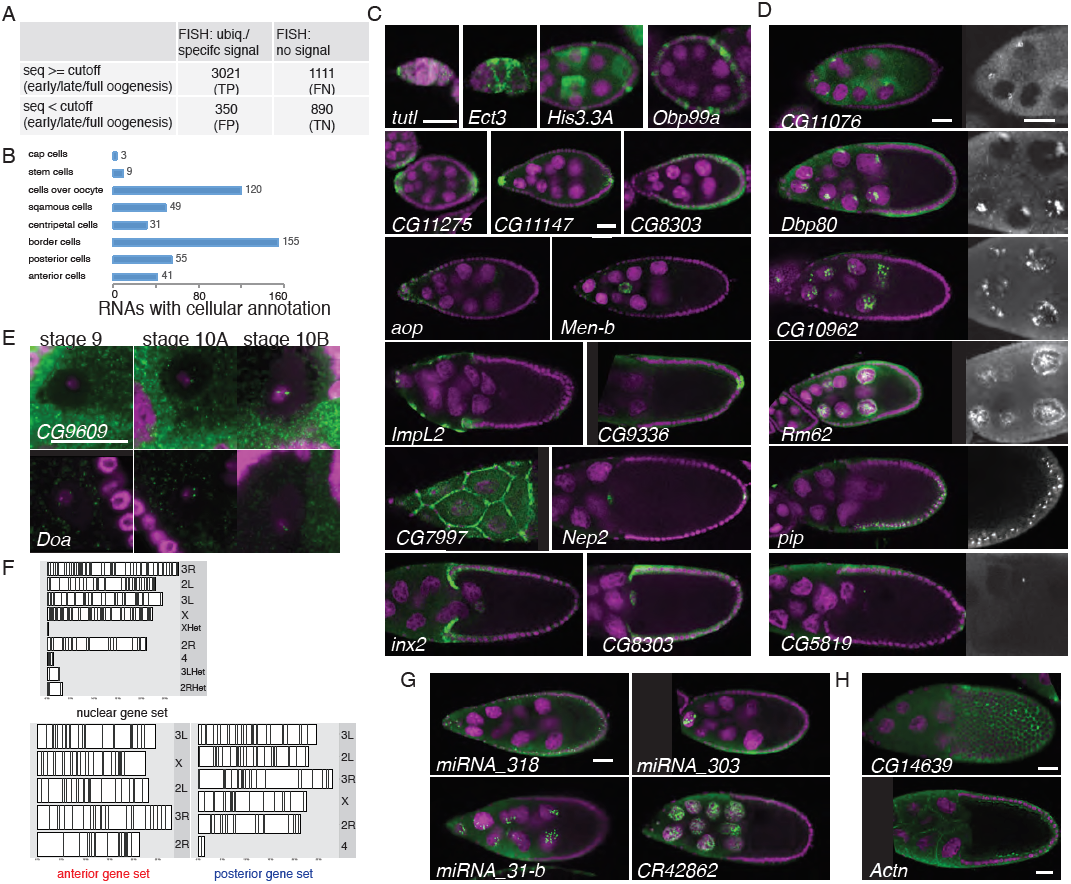
FISH screen results and controls. **(A)**. Estimate of false-positive/negative rate of the in situ screen using comparison with the independent transcriptomics data. A gene was classified as falsely positive if it was annotated as ubiquitous or specific by FISH but was not detectable by either 3Pseq or RNAseq at any time-point of oogenesis. In 20% of the experiments we failed to detect in situ signal (“no signal”) although the transcript was detected at least at one time point by at least one deep sequencing method. These may represent false negative results, possibly due to non-functional RNA probes, however we nevertheless included them in the downstream analysis in the no signal category (Figure S7). **(B)**. The cellular gene set was subcategorized according to the specific cellular expression pattern. Individual mRNAs can fall into several of these subgroups. **(C-D)**. Exemplary FISH experiments for the cellular **(C)** and nuclear **(D)** expression sets. RNA is shown in green and the DNA (labelled with DAPI) is shown in magenta. Scale bars: 30μm. *tutl* is expressed in cap cells at the tip of the germarium, while *Ect3* mRNA is detectable in the somatic epithelial cells of the germarium. Several mRNAs are expressed in mosaic pattern, indicating cell cycle control in somatic epithelial cells (*His3.3A, Obp99a*) and in nurse cells (*His3.3A*). Expression in the anterior and posterior follicle cells is often seen simultaneously (*CG11275, CG11147, Nep2*). Some mRNAs were expressed only in anterior follicle cells that become migratory border cells (*Men-b*) or in posterior follicle cells (*CG9336*). *CG8303* is expressed in the somatic cells destined to become columnar epithelium. aop is exclusively seen in follicle cells that will give rise to the squamous epithelium and several mRNAs are specifically expressed here at later stages (*ImpL2, CG7997*). mRNAs are also expressed in cells forming the border of columnar and squamous epithelial cells (*inx2*). Nuclei enrichments of RNAs in nurse cells can vary from a ring-like expression (*CG11076*), foci in a discrete area (*Dbp80*) to widespread foci (*CG10962*) or nucleoplasm signal (*Rm62*). RNAs are also detectable in epithelial cell nuclei (*pip*) and for 28 RNAs also in the oocyte nucleus (e.g. *CG5819*). Grey-scale image shows the respective RNA staining only in a zoomed-in view. **(E)**. *CG9609* and *Doa* mRNAs detected in the oocyte nucleus showing the enrichment over time at stages 9, 10A and 10B. At stage 9 only few small mRNA foci are visible, at stage 10 the mRNAs were enriched in proximity of the DNA in two large foci. **(F)**. Karyogram showing the chromosomal position of genes for nuclear, anterior and posterior localization classes. Neither nuclear RNA genes, which often appear in foci-like enrichments, nor anterior or posterior class genes are clustered on the chromosome. **(G)**. Examples of FISH experiment detecting distributions of non-coding RNA (in green). While *pri-miRNA-318* is enriched in somatic epithelial cell nuclei, *pri-miRNA-303,pri-miRNA-31-b* and the long non-coding RNA *CR42862* are restricted to nuclei of the germline nurse cells. Scale bar 30μm, DNA in magenta. **(H)**. mRNA enrichments in the somatic epithelial cells overlaying the oocyte (*CG14639*) and at the cortex of nurse cells (*Actn*). RNA signal shown in green. DNA, labelled with DAPI, is shown in magenta. Scale bar 30μm

**Figure 1-figure supplement 3.**
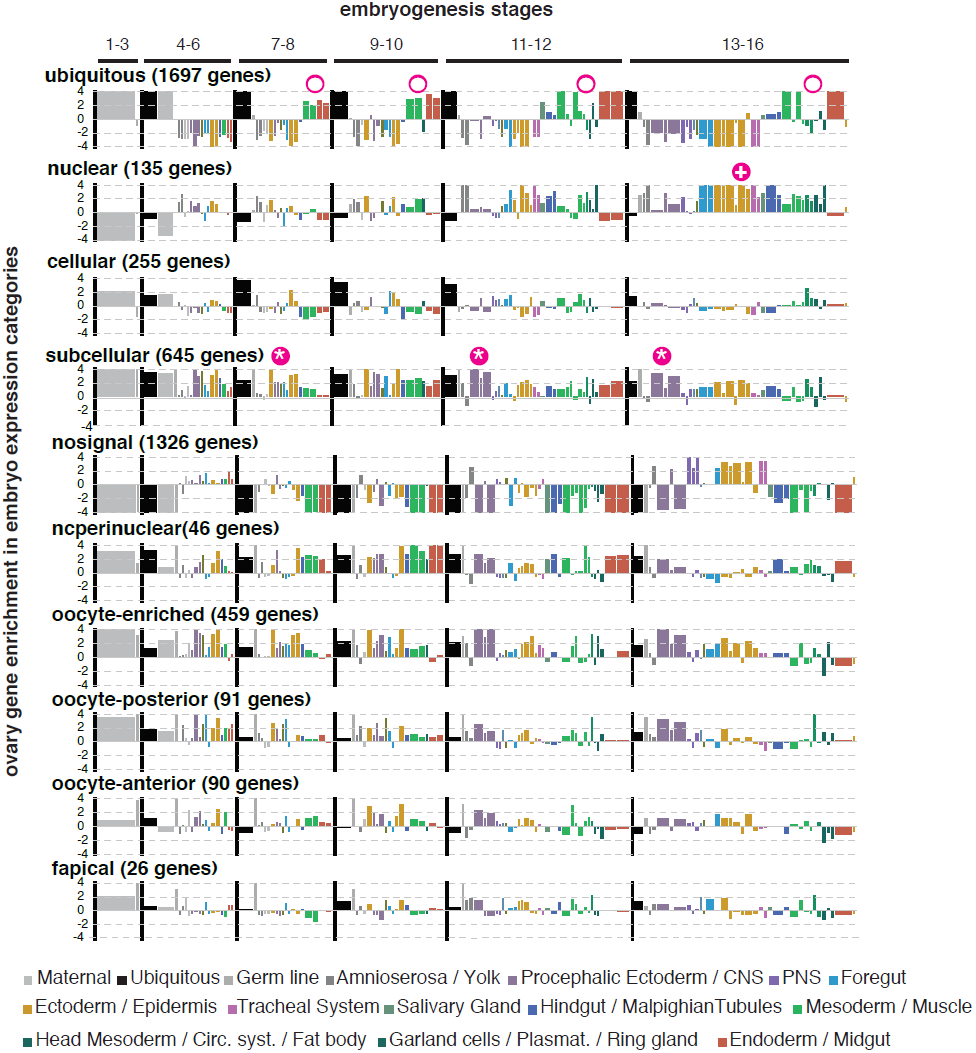
Ovary gene sets have specific expression patterns during embryogenesis. Linear hierarchy (Tomancak et al. 2007) plot showing at which embryonic stage and in which tissue the oogenesis gene sets re-expressed during embryogenesis. Each color-coded bar represents organ systems of the embryo from its stage specific anlagen to primordia to final differentiated structures. The width of the bar is proportional to the frequency with which this annotation term was used in the embryo dataset; the height corresponds to a z-score of over-(above axis) or under-representation (below axis) of the term in the set of genes defined by ovary annotation. The following oogenesis gene sets are shown: ubiquitous, nuclear, cellular, subcellular, no signal, nurse cells perinuclear, oocyte-enriched, oocyte anterior and oocyte posterior. Genes expressed ubiquitously in the ovary mostly remained ubiquitous in the embryo and were additionally enriched in meso-and endoderm (circle); Genes of the cellular gene set are enriched in ectoderm/epidermis cells of the late embryo (plus); subcellular genes were highly expressed in the ectoderm and nervous system (star) of the embryo. Most “no signal” genes are also underrepresented in almost all stages and tissues of embryogenesis, apart from the PNS and ectodermal derivatives in the late stages of embryogenesis. Perinuclear enriched genes are highly expressed in meso-and endoderm tissues. Oocyte enriched, oocyte anterior and oocyte posterior genes are overall very similarly expressed during embryogenesis, being high in the polarized CNS and ectoderm tissues.

**Figure 2-figure supplement 1.**
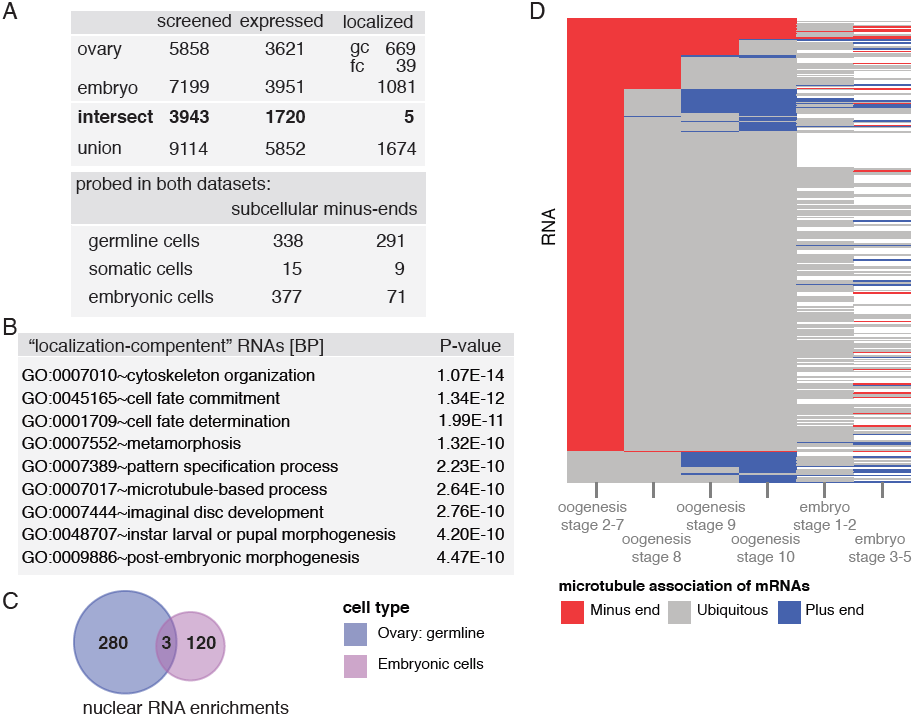
Changing localization of mRNAs in ovaries and embryos. **(A)**. Comparison of the results of FISH screens in ovaries and Drosophila embryos (Lecuyer et al., 2007). **(B)**. 1674 mRNAs show either during oogenesis or embryogenesis instances of subcellular localization (are “localization compe-tent”). These mRNAs are highly enriched for cytoskeletal/microtubule and cell fate/development biological functions. **(C)**. Venn diagram of the mRNAs showing nuclear enrichment in either oogenesis or embryogenesis. Only three mRNAs are nuclear at both developmental time-points. **(D)**. Expanded dendrogram from Figure 2b including the data for the first two time-points of embryogenesis (Lecuyer et al., 2007), showing that both microtubule minus-end (anterior) and micro-tubule plus-end (posterior) localization decreases with the onset of embryogenesis. Increase in minus-end localization is again observed at stage 3–5 of embryogenesis, when zygotic transcription is activated.

**Figure 2-figure supplement 2.**
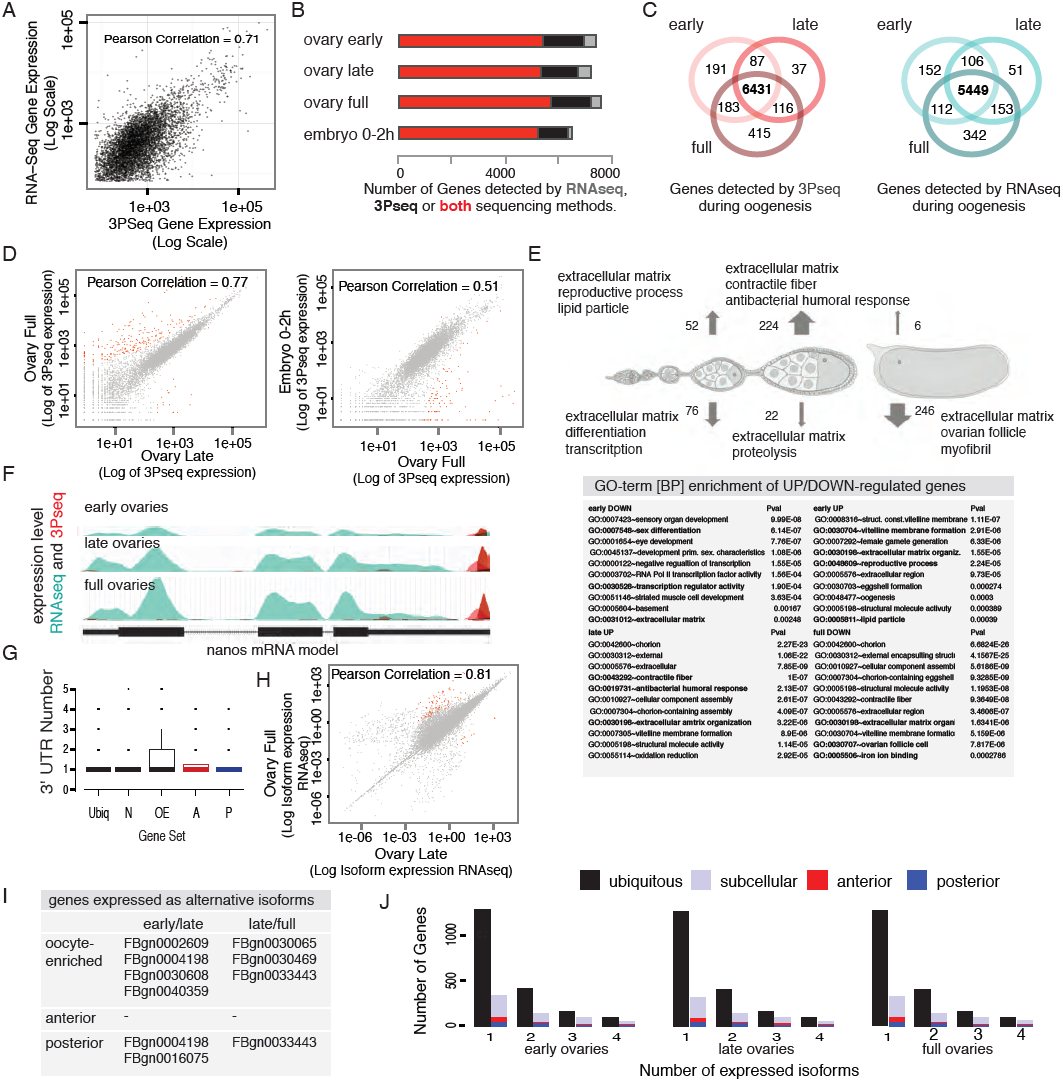
The transcriptome shows little variation over the course of oogenesis. **(A)**. Scatterplot of RNAseq and 3Pseq gene expression showing correlation (Pearson Correlation 0.71) between these two RNA sequencing methods. **(B)**. Results of transcriptome-wide sequencing from stage specific oogenesis samples (stage 1–7 = early, stage 9–10 = late, full ovaries) and 0–2 hour embryos. Across oogenesis and early embryogenesis, ∼5500 genes (red) were detected by both mRNA sequencing (RNAseq) and 3’prime end sequencing (3Pseq) at each stage while additional 1–2000 mRNAs were only captures with either RNAseq (grey) or 3Pseq (black) technique. Across all time-points about half of the *D.melanogaster* genome was expressed. **(C)**. More than 85% of the genes were expressed at each time point of oogenesis as shown by a Venn diagram overlapping the early, late and full ovarian transcriptome determined by 3Pseq (red) and RNAseq (turquois). **(D)**. Pair-wise correlation of late/full ovary datasets revealed that the stage-specific transcriptomes were highly similar (Pearson Correlation: 0.77); Significantly up-or down-regulated genes are shown in red (p-value adjusted for multiple testing < 0. 1). **(E)**. GO-term analysis of the genes identified as up (arrow up) and down (arrow down) regulated during oogenesis/early embryogenesis revealed that particularly genes encoding components of the extra-embryonic layers (vitelline membrane, ECM, cuticle) changed their expression levels. Late ovaries down-regulated genes and full ovaries up-regulated genes were not analysed since they contained too few entries. **(F)**. Example of the germline expressed *nanos* mRNA that shows a change in gene expression from early to full ovaries measured by RNAseq (green) and 3Pseq (red). The bottom part shows the nanos gene model with the position of introns and exons. **(G)**. Boxplot showing that the vast majority of genes express only one 3’UTR form during oogenesis, suggesting low prevalence of alternative poly-adenylation. **(H)**. Correlation analysis of expressed transcript isoform (deduced from RNAseq data) revealed that from early to late and from late to full ovaries almost no transcript-isoforms significantly changed in their expression level. Transcripts with significant changes are shown in red. **(I)**. Among the genes showing alternative isoform expression during oogenesis (see Figure 3D), few are found among subcellular enriched mRNAs, for example as oocyte enriched and posteriorly localized RNAs. No mRNA localized at the anterior pole exhibited alternative isoform expression. **(J)**. Number of transcripts per gene for the ubiquitous and subcellular gene set; Highlighted in red and blue are the anterior and posterior localized among the subcellular genes. The prevalence of alternatively spliced mRNAs is not changing between early, late and full ovary samples.

**Figure 3-figure supplement 1.**
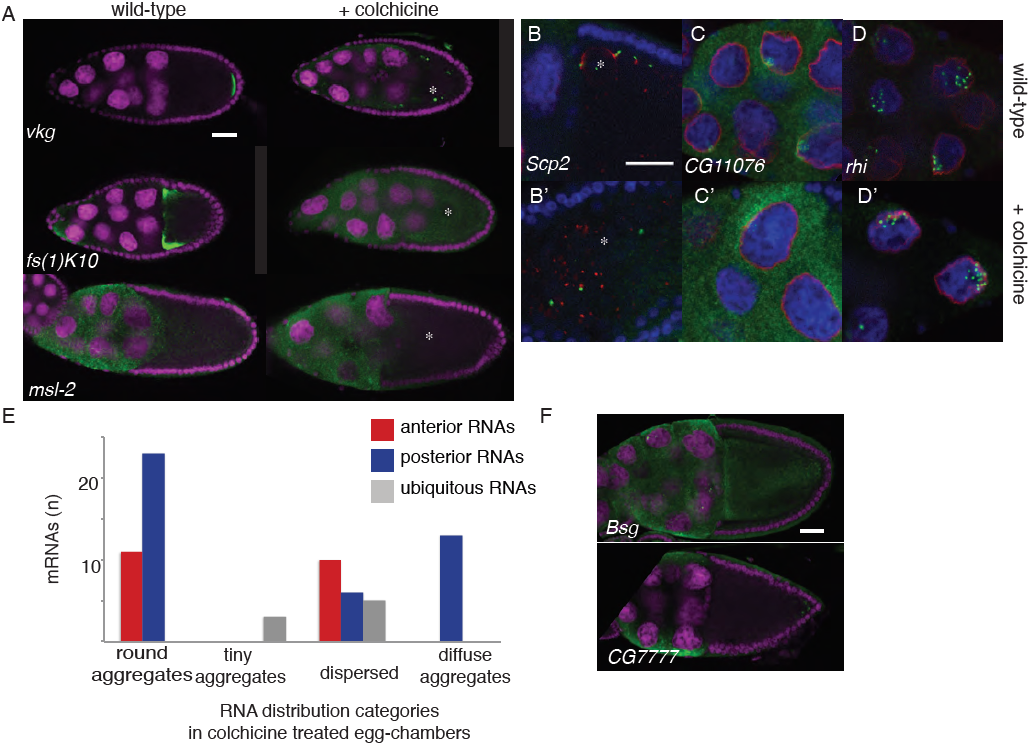
Cytoplasmic but not nuclear mRNA localization requires the cytoskeleton. **(A-D)**. Localization of the ubiquitous mRNA *msl-2* is unchanged upon microtubule depolymerization by colchicine, while another exemplary anterior and posterior mRNAa (*fs(1)K10, vkg*) become delocalized. mRNA localization in proximity to the nucleus is lost upon in colchicine treated egg-chambers (*Scp2*), RNAs localized partially nuclear and partially perinuclear loose the cytoplasmic localization (*CG11076*) while strictly nuclear RNAs are unaffected by microtubule depolymerization (*rhi*). **(E)**. Summary of the quality of anterior and posterior mRNA distributions upon microtubule depolymerization (round aggregates, tiny aggregates, dispersed and diffuse aggregates). Diffuse aggre-gates were observed for those mRNAs that in wild type egg-chambers showed a diffuse posterior enrichment (e.g. Figure 1b: *fs(1)N*). **(F)**. *Bsg* and *CG7777* mRNA distribution is impaired in in egg-chambers lacking posterior *oskar* mRNA. **(A-F)**. FISH experiments showing the RNA in in green; DNA (labelled with DAPI) is shown in magenta **(A, F)** or blue **(B-D)** and the nuclear membrane is stained with WGA dye shown in red **(B-D)**. Scale bar 30μm.

**Figure 4-figure supplement 1.**
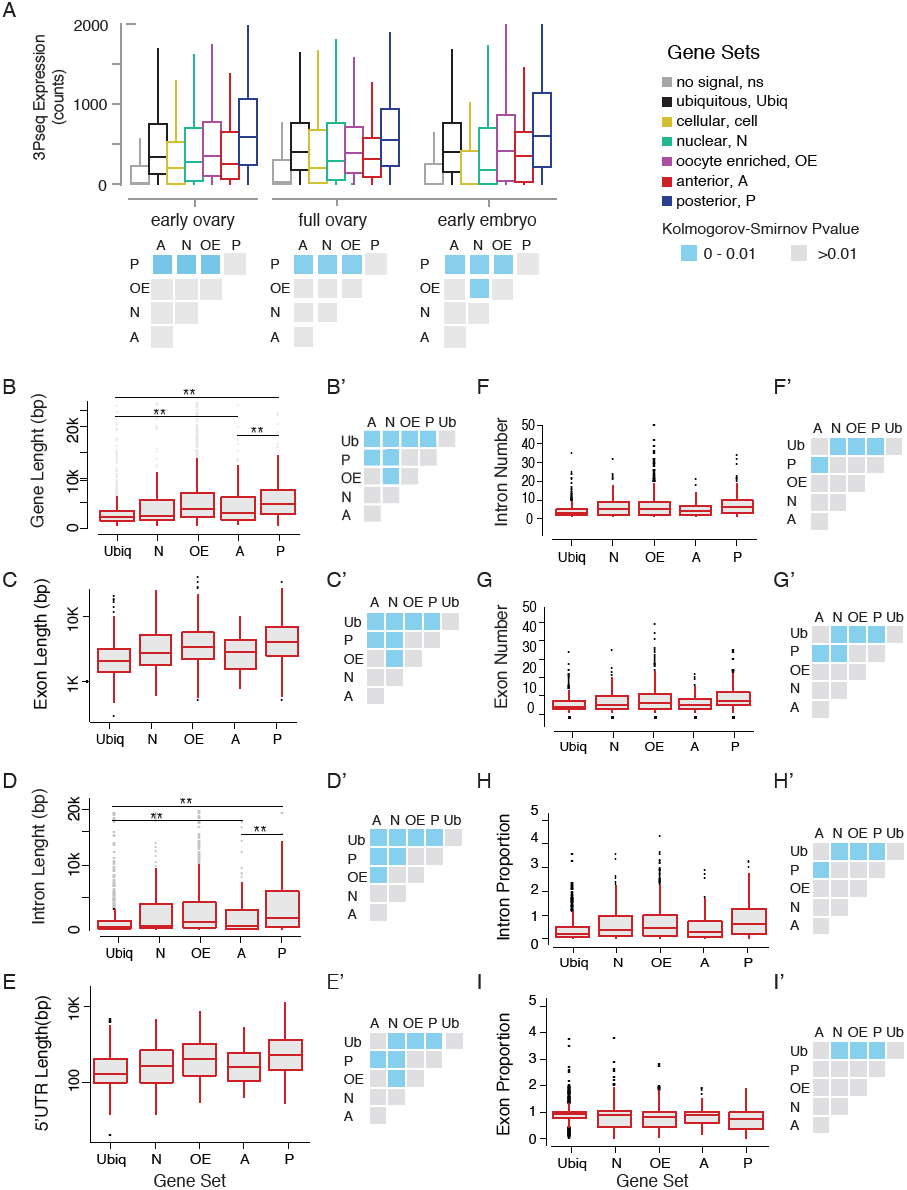
Gene Features of subcellular enriched mRNAs. **(A)**. Boxplots showing the median mRNA expression measured by 3Pseq per gene set in early and full ovaries and in 0–2h embryos. At the onset of embryogenesis, the cellular mRNAs were almost as low as the “no signal” class, confirming their predominant expression in somatic cells that at this time-point have undergone apoptosis. Accompanying the boxplot is the matrix of statistical significance tests (Kolmogorov-Smirnov) of the null hypothesis that the distributions of median expression values across the subcellular gene sets are the same. Statistically significant differences (p-value <0.01) are shown in blue, while gene sets that did not differ significantly are shown in grey (p-value >0.01). **(B-I)**. Boxplots showing the median gene length **(B)**, exon length **(C)**, intron length **(D)**, 5’UTR length **(E)**, intron number **(F)**, exon number **(G)**, intron proportion **(H)** and exon proportion **(I)** for each gene set and the corresponding significance level calculated by the Kolmogorov-Smirnov (KS) test. Statistically significant differences (p-value <0.01) are shown in blue, while gene sets that did not differ significantly are shown 31 in grey (p-value >0.01).

## References

Amrute-Nayak, M. & Bullock, S. L. 2012. Single-molecule assays reveal that RNA localization signals regulate dynein-dynactin copy number on individual transcript cargoes. Nature cell biology, 14, 416–23, 10.1038/ncb2446.

Anders, S. & Huber, W. 2010. Differential expression analysis for sequence count data. Genome biology, 11, R106, 10.1186/gb-2010-11-10-r106.

Ashton-Beaucage, D., Udell, C. M., Lavoie, H., Baril, C., Lefrancois, M., Chagnon, P., Gendron, P., Caron-Lizotte, O., Bonneil, E., Thibault, P. & Therrien, M. 2010. The exon junction complex controls the splicing of MAPK and other long intron-containing transcripts in Drosophila. Cell, 143, 251–62, 10.1016/j.cell.2010.09.014.

Berleth, T., Burri, M., Thoma, G., Bopp, D., Richstein, S., Frigerio, G., Noll, M. & Nusslein-Volhard, C. 1988. The role of localization of bicoid RNA in organizing the anterior pattern of the Drosophila embryo. EMBO J, 7, 1749–56.

Besse, F., Lopez De Quinto, S., Marchand, V., Trucco, A. & Ephrussi, A. 2009. Drosophila PTB promotes formation of high-order RNP particles and represses oskar translation. Genes & development, 23, 195–207, 10.1101/gad.505709.

Bullock, S. L. 2011. Messengers, motors and mysteries: sorting of eukaryotic mRNAs by cytoskeletal transport. Biochemical Society transactions, 39, 1161–5, 10.1042/BST0391161.

Bullock, S. L. & Ish-Horowicz, D. 2001. Conserved signals and machinery for RNA transport in Drosophila oogenesis and embryogenesis. Nature, 414, 611–6, 10.1038/414611a.

Bullock, S. L., Ringel, I., Ish-Horowicz, D. & Lukavsky, P. J. 2010. A’-form RNA helices are required for cytoplasmic mRNA transport in Drosophila. Nature structural & molecular biology, 17, 703–9, 10.1038/nsmb.1813.

Calderone, A., Castagnoli, L. & Cesareni, G. 2013. mentha: a resource for browsing integrated protein-interaction networks. Nature methods, 10, 690–1, 10.1038/nmeth.2561.

Cha, B. J., Serbus, L. R., Koppetsch, B. S. & Theurkauf, W. E. 2002. Kinesin I-dependent cortical exclusion restricts pole plasm to the oocyte posterior. Nat Cell Biol, 4, 592–8, 10.1038/ncb832.

Chao, J. A., Patskovsky, Y., Patel, V., Levy, M., Almo, S. C. & Singer, R. H. 2010. ZBP1 recognition of beta-actin zipcode induces RNA looping. Genes & development, 24, 148–58, 10.1101/gad.1862910.

Chintapalli, V. R., Wang, J. & Dow, J. A. 2007. Using FlyAtlas to identify better Drosophila melanogaster models of human disease. Nat Genet, 39, 715–20, 10.1038/ng2049.

Dienstbier, M., Boehl, F., Li, X. & Bullock, S. L. 2009. Egalitarian is a selective RNA-binding protein linking mRNA localization signals to the dynein motor. Genes & development, 23, 1546–58, 10.1101/gad.531009.

Dix, C. I., Soundararajan, H. C., Dzhindzhev, N. S., Begum, F., Suter, B., Ohkura, H., Stephens, E. & Bullock, S. L. 2013. Lissencephaly-1 promotes the recruitment of dynein and dynactin to transported mRNAs. The Journal of cell biology, 202, 479–94, 10.1083/jcb.201211052.

Ephrussi, A., Dickinson, L. K. & Lehmann, R. 1991. Oskar organizes the germ plasm and directs localization of the posterior determinant nanos. Cell, 66, 37–50.

Forrest, K. M. & Gavis, E. R. 2003. Live imaging of endogenous RNA reveals a diffusion and entrapment mechanism for nanos mRNA localization in Drosophila. Curr Biol, 13, 1159–68.

Ghosh, S., Marchand, V., Gaspar, I. & Ephrussi, A. 2012. Control of RNP motility and localization by a splicing-dependent structure in oskar mRNA. Nature structural & molecular biology, 19, 441–9, 10.1038/nsmb.2257.

Graveley, B. R., Brooks, A. N., Carlson, J. W., Duff, M. O., Landolin, J. M., Yang, L., Artieri, C. G., Van Baren, M. J., Boley, N., Booth, B. W., Brown, J. B., Cherbas, L., Davis, C. A., Dobin, A., Li, R., Lin, W., Malone, J. H., Mattiuzzo, N. R., Miller, D., Sturgill, D., Tuch, B. B., Zaleski, C., Zhang, D., Blanchette, M., Dudoit, S., Eads, B., Green, R. E., Hammonds, A., Jiang, L., Kapranov, P., Langton, L., Perrimon, N., Sandler, J. E., Wan, K. H., Willingham, A., Zhang, Y., Zou, Y., Andrews, J., Bickel, P. J., Brenner, S. E., Brent, M. R., Cherbas, P., Gingeras, T. R., Hoskins, R. A., Kaufman, T. C., Oliver, B. & Celniker, S. E. 2011. The developmental transcriptome of Drosophila melanogaster. Nature, 471, 473–9, 10.1038/nature09715.

Hachet, O. & Ephrussi, A. 2004. Splicing of oskar RNA in the nucleus is coupled to its cytoplasmic localization. Nature, 428, 959–63, 10.1038/nature02521.

Horne-Badovinac, S. & Bilder, D. 2008. Dynein regulates epithelial polarity and the apical localization of stardust A mRNA. PLoS genetics, 4, e8, 10.1371/journal.pgen.0040008.

Huang Da, W., Sherman, B. T. & Lempicki, R. A. 2009. Systematic and integrative analysis of large gene lists using DAVID bioinformatics resources. Nature protocols, 4, 44–57, 10.1038/nprot.2008.211.

Hubisz, M. J., Pollard, K. S. & Siepel, A. 2011. PHAST and RPHAST: phylogenetic analysis with space/time models. Briefings in bioinformatics, 12, 41–51, 10.1093/bib/bbq072.

Jambhekar, A. & Derisi, J. L. 2007. Cis-acting determinants of asymmetric, cytoplasmic RNA transport. RNA, 13, 625–42, 10.1261/rna.262607.

Jambor, H., Brunel, C. & Ephrussi, A. 2011. Dimerization of oskar 3’ UTRs promotes hitchhiking for RNA localization in the Drosophila oocyte. RNA, 17, 2049–57, 10.1261/rna.2686411.

Jambor, H., Mueller, S., Bullock, S. L. & Ephrussi, A. 2014. A stem-loop structure directs oskar mRNA to microtubule minus ends. RNA, 20, 429–39, 10.1261/rna.041566.113.

Januschke, J., Gervais, L., Gillet, L., Keryer, G., Bornens, M. & Guichet, A. 2006. The centrosome-nucleus complex and microtubule organization in the Drosophila oocyte. Development, 133, 129–39, 10.1242/dev.02179.

Jenny, A., Hachet, O., Zavorszky, P., Cyrklaff, A., Weston, M. D., Johnston, D. S., Erdelyi, M. & Ephrussi, A. 2006. A translation-independent role of oskar RNA in early Drosophila oogenesis. Development, 133, 2827–33, 10.1242/dev.02456.

Kislauskis, E. H., Zhu, X. & Singer, R. H. 1994. Sequences responsible for intracellular localization of beta-actin messenger RNA also affect cell phenotype. J Cell Biol, 127, 441–51.

Lange, S., Katayama, Y., Schmid, M., Burkacky, O., Brauchle, C., Lamb, D. C. & Jansen, R. P. 2008. Simultaneous transport of different localized mRNA species revealed by live-cell imaging. Traffic, 9, 1256–67, 10.1111/j.1600–0854.2008.00763.x.

Lecuyer, E., Yoshida, H., Parthasarathy, N., Alm, C., Babak, T., Cerovina, T., Hughes, T. R., Tomancak, P. & Krause, H. M. 2007. Global analysis of mRNA localization reveals a prominent role in organizing cellular architecture and function. Cell, 131, 174–87, 10.1016/j.cell.2007.08.003.

Lehmann, R. & Nusslein-Volhard, C. 1986. Abdominal segmentation, pole cell formation, and embryonic polarity require the localized activity of oskar, a maternal gene in Drosophila. Cell, 47, 141–52.

Manseau, L. J. & Schupbach, T. 1989. cappuccino and spire: two unique maternal-effect loci required for both the anteroposterior and dorsoventral patterns of the Drosophila embryo. Genes Dev, 3, 1437–52.

Medioni, C., Mowry, K. & Besse, F. 2012. Principles and roles of mRNA localization in animal development. Development, 139, 3263–76, 10.1242/dev.078626.

Micklem, D. R., Adams, J., Grunert, S. & St Johnston, D. 2000. Distinct roles of two conserved Staufen domains in oskar mRNA localization and translation. EMBO J, 19, 1366–77, 10.1093/emboj/19.6.1366.

Mikl, M., Vendra, G. & Kiebler, M. A. 2011. Independent localization of MAP2, CaMKIIalpha and beta-actin RNAs in low copy numbers. EMBO reports, 12, 1077–84, 10.1038/embor.2011.149.

Neuman-Silberberg, F. S. & Schüpbach, T. 1993. The Drosophila dorsoventral patterning gene gurken produces a dorsally localized RNA and encodes a TGFa-like protein. Cell, 75, 165–174.

Park, H. Y., Lim, H., Yoon, Y. J., Follenzi, A., Nwokafor, C., Lopez-Jones, M., Meng, X. & Singer, R. H. 2014. Visualization of dynamics of single endogenous mRNA labeled in live mouse. Science, 343, 422–4, 10.1126/science.1239200.

Pollard, K. S., Hubisz, M. J., Rosenbloom, K. R. & Siepel, A. 2010. Detection of nonneutral substitution rates on mammalian phylogenies. Genome research, 20, 110–21, 10.1101/gr.097857.109.

Preitner, N., Quan, J., Nowakowski, D. W., Hancock, M. L., Shi, J., Tcherkezian, J., Young-Pearse, T. L. & Flanagan, J. G. 2014. APC is an RNA-binding protein, and its interactome provides a link to neural development and microtubule assembly. Cell, 158, 368–82, 10.1016/j.cell.2014.05.042.

Roth, S., Neuman-Silberberg, F. S., Barcelo, G. & Schupbach, T. 1995. cornichon and the EGF receptor signaling process are necessary for both anterior-posterior and dorsal-ventral pattern formation in Drosophila. Cell, 81, 967–78.

Saunders, C. & Cohen, R. S. 1999. The role of oocyte transcription, the 5’UTR, and translation repression and derepression in Drosophila gurken mRNA and protein localization. Mol Cell, 3, 43–54.

Schindelin, J., Arganda-Carreras, I., Frise, E., Kaynig, V., Longair, M., Pietzsch, T., Preibisch, S., Rueden, C., Saalfeld, S., Schmid, B., Tinevez, J. Y., White, D. J., Hartenstein, V., Eliceiri, K., Tomancak, P. & Cardona, A. 2012. Fiji: an open-source platform for biological-image analysis. Nature methods, 9, 676–82, 10.1038/nmeth.2019.

Sinsimer, K. S., Jain, R. A., Chatterjee, S. & Gavis, E. R. 2011. A late phase of germ plasm accumulation during Drosophila oogenesis requires lost and rumpelstiltskin. Development, 138, 3431–40, 10.1242/dev.065029.

Snee, M. J., Arn, E. A., Bullock, S. L. & Macdonald, P. M. 2005. Recognition of the bcd mRNA localization signal in Drosophila embryos and ovaries. Molecular and cellular biology, 25, 1501–10, 10.1128/MCB.25.4.1501-1510.2005.

Soundararajan, H. C. & Bullock, S. L. 2014. The influence of dynein processivity control, MAPs, and microtubule ends on directional movement of a localising mRNA. eLife, 3, e01596, 10.7554/eLife.01596.

St Johnston, D., Beuchle, D. & Nusslein-Volhard, C. 1991. Staufen, a gene required to localize maternal RNAs in the Drosophila egg. Cell, 66, 51–63.

St Johnston, D., Driever, W., Berleth, T., Richstein, S. & Nusslein-Volhard, C. 1989. Multiple steps in the localization of bicoid RNA to the anterior pole of the Drosophila oocyte. Development, 107 Suppl, 13–9.

Tanaka, T. & Nakamura, A. 2008. The endocytic pathway acts downstream of Oskar in Drosophila germ plasm assembly. Development, 135, 1107–17.

Theurkauf, W. E., Smiley, S., Wong, M. L. & Alberts, B. M. 1992. Reorganization of the cytoskeleton during Drosophila oogenesis: implications for axis specification and intercellular transport. Development, 115, 923–36.

Tomancak, P., Berman, B. P., Beaton, A., Weiszmann, R., Kwan, E., Hartenstein, V., Celniker, S. E. & Rubin, G. M. 2007. Global analysis of patterns of gene expression during Drosophila embryogenesis. Genome Biol, 8, R145, gb-2007-8-7-r145 [pii]10.1186/gb-2007-8-7-r145.

Ulitsky, I., Shkumatava, A., Jan, C. H., Subtelny, A. O., Koppstein, D., Bell, G. W., Sive, H. & Bartel, D. P. 2012. Extensive alternative polyadenylation during zebrafish development. Genome research, 22, 2054–66, 10.1101/gr.139733.112.

Van Eeden, F. J., Palacios, I. M., Petronczki, M., Weston, M. J. & St Johnston, D. 2001. Barentsz is essential for the posterior localization of oskar mRNA and colocalizes with it to the posterior pole. J Cell Biol, 154, 511–24.

Vanzo, N., Oprins, A., Xanthakis, D., Ephrussi, A. & Rabouille, C. 2007. Stimulation of endocytosis and actin dynamics by Oskar polarizes the Drosophila oocyte. Dev Cell, 12, 543–55, 10.1016/j.devcel.2007.03.002.

Vanzo, N. F. & Ephrussi, A. 2002. Oskar anchoring restricts pole plasm formation to the posterior of the Drosophila oocyte. Development, 129, 3705–14.

Whittaker, K. L., Ding, D., Fisher, W. W. & Lipshitz, H. D. 1999. Different 3’ untranslated regions target alternatively processed hu-li tai shao (hts) transcripts to distinct cytoplasmic locations during Drosophila oogenesis. J Cell Sci, 112 (Pt 19), 3385–98.

Zimyanin, V., Lowe, N. & St Johnston, D. 2007. An oskar-dependent positive feedback loop maintains the polarity of the Drosophila oocyte. Current biology : CB, 17, 353–9, 10.1016/j.cub.2006.12.044.

Zimyanin, V. L., Belaya, K., Pecreaux, J., Gilchrist, M. J., Clark, A., Davis, I. & St Johnston, D. 2008. In vivo imaging of oskar mRNA transport reveals the mechanism of posterior localization. Cell, 134, 843–53, 10.1016/j.cell.2008.06.053.

